# Evolution, diversity, function, and marker-based elimination of the disease susceptibility gene *Snn1* in wheat

**DOI:** 10.1101/2024.01.24.577084

**Authors:** Sudeshi Seneviratne, Gongjun Shi, Agnes Szabo-Hever, Zengcui Zhang, Amanda R. Peters Haugrud, Katherine L. D. Running, Gurminder Singh, Raja Sekhar Nandety, Jason D. Fiedler, Phillip E. McClean, Steven S. Xu, Timothy L. Friesen, Justin D. Faris

## Abstract

Septoria nodorum blotch (SNB), caused by *Parastagonospora nodorum,* is a disease of durum and common wheat initiated by the recognition of pathogen-produced necrotrophic effectors (NEs) by specific wheat genes. The wheat gene *Snn1* encodes a wall-associated kinase that directly interacts with the NE SnTox1 leading to the development of SNB. Here, sequence analysis of *Snn1* from 114 accessions including diploid, tetraploid and hexaploid wheat species revealed that some wheat lines possess two copies of *Snn1* (designated *Snn1-B1* and *Snn1-B2*) approximately 120 kb apart. *Snn1-B2* evolved relatively recently as a paralog of *Snn1-B1*, and both genes have undergone diversifying selection. Three point mutations associated with the formation of the first SnTox1-sensitive *Snn1-B1* allele from a primitive wild wheat were identified. Four subsequent and independent SNPs, three in *Snn1-B1* and one in *Snn1-B2*, converted the sensitive alleles to insensitive forms. Protein modeling indicated these four mutations could abolish *Snn1*-SnTox1 compatibility either through destabilization of the Snn1 protein or direct disruption of the protein-protein interaction. High-throughput markers were developed for the causal mutations and evaluated on panels of durum and common wheat. The markers were able to correctly identify 96.9 % of SnTox1-sensitive durum wheat accessions, and a marker for the null allele was 100% accurate at predicting SnTox1-insensitive lines in both durum and spring wheat. Results of this study increase our understanding of the evolution, diversity, and function of *Snn1-B1* and *Snn1-B2* genes and will be useful for marker-assisted elimination of these genes for better host resistance.

## Introduction

With an annual global production of ∼ 770 million tons (FAO 2022), common wheat (*Triticum aestivum* L., 2*n* = 6*x* = 42, AABBDD) and durum wheat (*T. turgidum* ssp*. durum* (Desf.) Husnot., 2*n* = 4*x* = 28, AABB) serve as major sources of sustenance throughout the world. Therefore, factors that affect wheat yield and quality have a significant impact on the agricultural economy worldwide. Necrotrophic fungal pathogens are a major biotic threat to global wheat production, and studies have indicated that global climate change could promote the colonization of necrotrophs in plants at an accelerated rate (Manning et al. 1995). Therefore, it is important to improve our knowledge of the interactions between plants and necrotrophic pathogens.

Necrotrophs require dead or dying tissue on which to feed, and studies have shown that some necrotrophs have acquired the ability to obtain nutrients from plants by hijacking the defense mechanisms that plants have evolved to fight biotrophic pathogens(reviewed by Faris and Friesen 2020; Friesen and Faris 2010, 2021). These pathogens produce necrotrophic effectors (NEs) that are recognized by specific dominant host genes in an inverse gene-for-gene manner to trigger responses that lead to death of plant tissue (i.e. host-induced programmed cell death) creating an environment favorable to the necrotroph’s lifestyle, which would otherwise be unfavorable to the survival of a biotrophic pathogen. In these systems, the presence of a pathogen-produced NE and the corresponding dominant host gene for sensitivity leads to a compatible interaction, and ultimately, disease susceptibility. If either the NE or the dominant host allele are absent, a resistance response occurs.

Septoria nodorum blotch (SNB) is a severe foliar and glume disease that can cause yield losses as high as 30 % in wheat (reviewed by Downie et al. 2021). SNB has been reported in most wheat-growing areas including Australia, South Asia, North Africa, Europe, and North America (Oliver et al. 2012; Bearchell et al. 2005; Crook et al. 2012). The disease is caused by *Parastagonospora* [syn. anamorph: *Stagonospora*; teleomorph: *Phaeosphaeria*] *nodorum*, which is a heterothallic, necrotrophic, filamentous, ascomycete fungus pathogenic on wheat, barley, and a wide range of wild grasses.

Multiple sensitivity genes that recognize NEs produced by *P. nodorum* have been identified (see Friesen and Faris 2021; Peters Haugrud et al. 2022 and citations within for review), and several of them (*Tsn1*, *Snn1*, *Snn3*-*D1, Snn3-B1, Snn3-B2, Snn5-B1*) have been cloned (Faris et al. 2010; Shi et al. 2016; Zhang et al. 2021; J. Faris, K. Running, Z. Zhang, unpublished). *Snn1*-SnTox1 was the first sensitivity gene-NE interaction identified, and *Snn1* was the second host sensitivity gene cloned in the wheat-*P. nodorum* pathosystem (Liu et al. 2004a; Liu et al. 2004b; Shi et al. 2016). *Snn1* encodes a protein that contains a signal sequence, a wall-associated receptor kinase galacturonan binding domain (GUB_WAK), an epidermal growth factor-calcium binding domain (EGF_CA), a transmembrane domain, and a serine/threonine protein kinase (S/TPK) domain making it a member of the wall-associated kinase (WAK) class of receptor kinase genes (Shi et al. 2016). A yeast two-hybrid assay revealed that the Snn1 and SnTox1 proteins interact directly, with SnTox1 binding to a region between the GUB_WAK and EGF_CA domains of Snn1 (Shi et al. 2016).

A compatible *Snn1*-SnTox1 interaction can play a major role in the development of SNB of wheat (Liu et al. 2004b). This interaction is important because SnTox1 was found to be present in 95.4% of *P. nodorum* isolates collected from spring, winter, and durum wheat-growing regions in the US (Richards et al. 2019), and it was present in 84% of a global collection of *P. nodorum* isolates (McDonald et al. 2013). However, the percentage of SnTox1-sensitive lines widely fluctuates between different germplasm groups worldwide (Peters Haugrud et al. 2022). Shi et al. (2016) reported that *Snn1* is more common among tetraploid wheat lines compared to hexaploids.

It is important to understand the evolutionary forces of selection to allow breeders to directly access key alleles that confer resistance. In the case of *Snn1*, resistance is conferred by the lack of a functional susceptibility allele, i.e. the functional *Snn1* allele must be removed or disrupted to eliminate recognition of the pathogen. In-depth analysis of the evolution and diversity of *Snn1* alleles should yield knowledge regarding causal polymorphisms within the *Snn1* gene that can be targeted for the development of functional markers for marker-assisted elimination (MAE) of functional *Snn1* alleles. Therefore, the primary objective of this research was to develop diagnostic markers suitable for high-throughput MAE of *Snn1* from breeding material. Toward that goal, we acquired knowledge and insights regarding the evolution, natural and induced variation, and functional aspects of *Snn1* and *Snn1*-SnTox1 interactions.

## Results

### Reaction of wheat lines to SnTox1

Screening of the wheat accessions with SnTox1 showed that the lines had different levels of sensitivity to SnTox1. An expanded scoring scale, which included seven categories (0, 0.5, 1.0, 1.5, 2.0, 2.5, and 3.0), was developed for rating SnTox1 sensitivity levels based on the one defined by Zhang et al. (2011) (Fig. 1). Lines with scores ≤ 1.0 were considered as insensitive and lines with scores > 1.0 were considered as sensitive.

**Figure 1.**
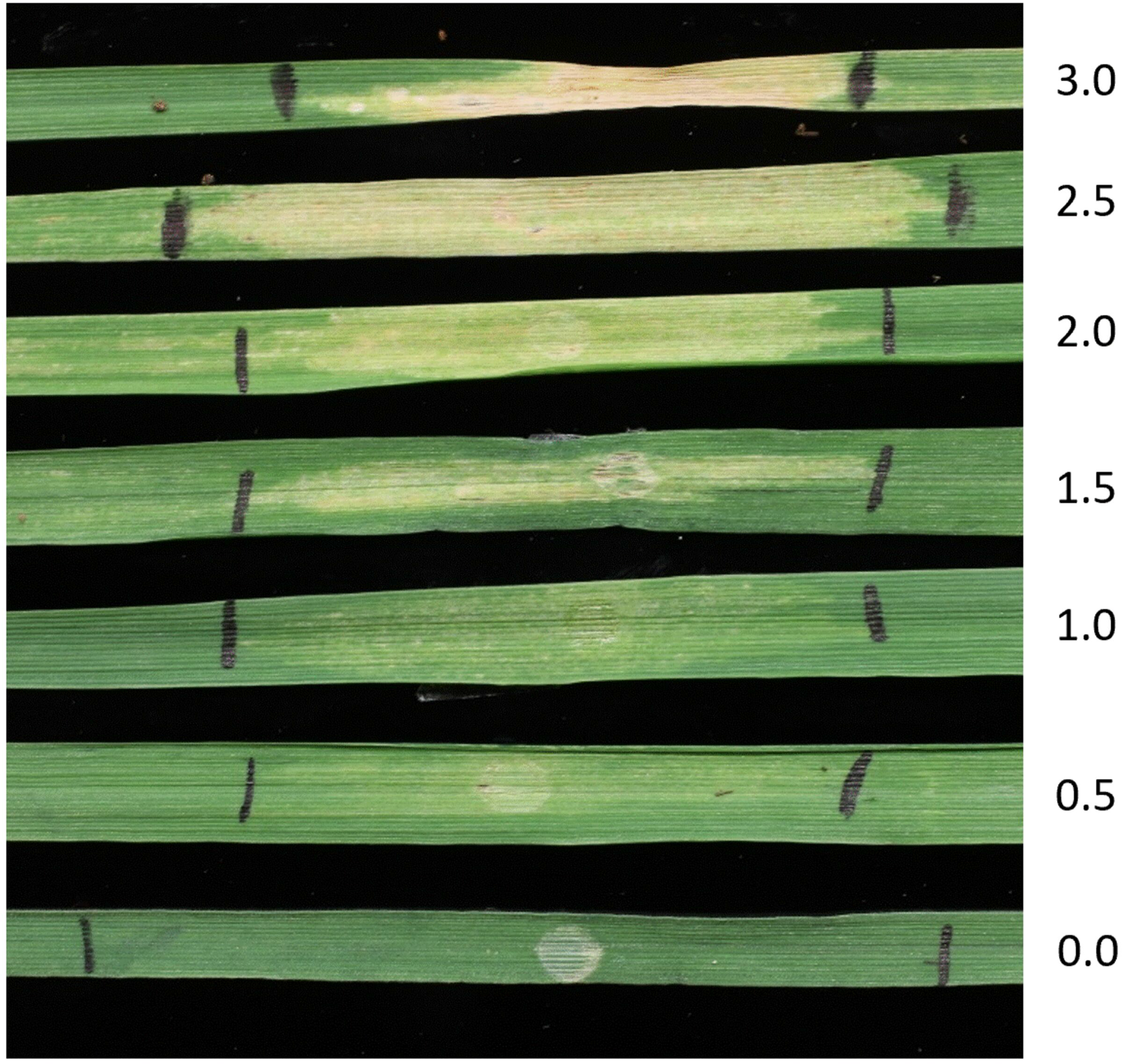
The scoring scale used for phenotyping wheat leaves after infiltration with SnTox1. A score of 0 indicates an insensitive response and a score of 3.0 indicates a severely sensitive response. Lines with scores ≤ 1.0 were considered as insensitive and lines with scores > 1.0 were considered as sensitive.

Bartlett’s Chi-squared test for homogeneity of variances showed that the variance among the replicates was not significantly different in any of the experiments (Supplemental Table S1). Therefore, the means from each experiment were used for further analysis.

### Analysis of *Snn1* copy number and expression

Sequencing of *Snn1* and analysis of the sequencing reads revealed that some lines contained double peaks at single bases in the chromatograms of sequence reads, indicating that two copies of the gene existed and were being amplified. Among the 114 lines from which we sequenced *Snn1* in this study, 92 had only one copy and 22 had two copies (Supplemental Table S2). The original copy was designated as *Snn1-B1*, and the second copy identified in this research was designated *Snn1-B2* (Fig. 2) in accordance with the guidelines for wheat gene nomenclature (Boden et al. 2023). Two SNPs that were polymorphic between *Snn1-B1* and *Snn1-B2* (G127A and A264G) in lines that had both copies were identified by sequence alignment. Both were located within the coding region of the gene (Fig. 2). G127A encoded a missense variant V43I, and A264G was a synonymous mutation. These two SNPs were used to differentiate the two copies where *Snn1-B1* had an A at position 127 and a G at position 264, whereas *Snn1-B2* had a G at position 127 and an A at position 264. However, four lines in which both copies had A at position 127 and G at 264 were observed (FHB4512, Langdon, Opata85, Renan), and the fact they contained two copies was revealed through sequencing. In FHB4512, three SNPs (T531C, C707T, A1556G) distinguished the two copies, whereas in Langdon, only one SNP (C2187A) differentiated them. Opata85 and Renan had more SNPs between the two copies, with Opata85 having a total of 26 SNPs, including C707T, and Renan having 37 SNPs, including C2187A. GenBank accession numbers of all the sequences can be found in Supplemental Tables S2 and S3. For the lines in which SNP G127A was absent, it could not be determined which copy was *Snn1-B1* and which was *Snn1-B1* relative to the described copies in other lines.

**Figure 2.**
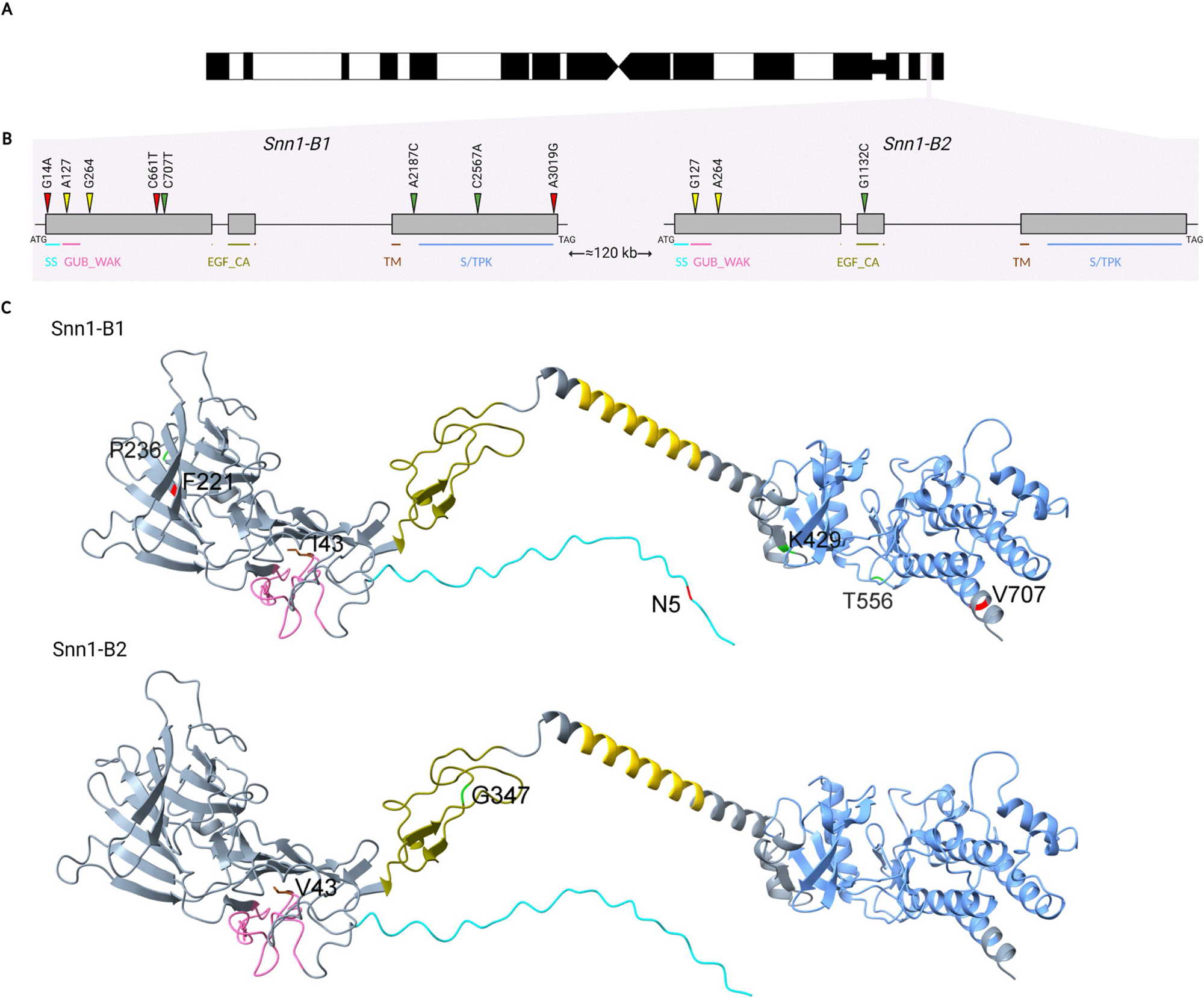
Illustration of the *Snn1* genes and their encoded proteins. A: Chromosome 1B with the position of the genomic region containing *Snn1* genes indicated on the short arm satellite. B: Gene structures and positions of major domains and critical mutations. The SNPs that differentiate *Snn1-B1* and *Snn1-B2* are shown in yellow. The mutations associated with the evolutionary change from insensitivity to sensitivity are shown in red and the mutations that change the phenotype from sensitive to insensitive are shown in green. SS: Signal sequence; GUB_WAK: Wall-associated receptor kinase galacturonan binding domain; EGF_CA: Epidermal growth factor-calcium binding domain; TM: Transmembrane domain; S/TPK: Serine/threonine protein kinase domain. C: Predicted structures of Snn1-B1 and Snn1-B2 proteins. The SS, GUB_WAK, EGF_CA, TM, and S/TPK domains are shown in cyan, pink, olive green, yellow, and blue, respectively. Critical amino acids associated with the change from insensitivity to sensitivity are shown in red (N5, F221, V707) and those associated with the change in phenotype from sensitive to insensitive are shown in green (P236, G347, K429, T556). The single amino acid that differentiates Snn1-B1 and Snn1-B2 is highlighted in brown (I43V). Created with BioRender.com.

Sequencing of cDNA revealed that when a single copy of the gene was present in the genome, it was expressed, but among lines that possessed both genes, only *Snn1-B1* was transcribed (Supplemental Fig. S5). Thirty-five lines expressing *Snn1-B1* and 57 lines expressing *Snn1-B2* were observed among lines with a single copy of *Snn1* (Supplemental Table S2).

To determine the relative physical positions of the two copies of *Snn1*, we evaluated the available wheat genome pseudochromosome assemblies with BLASTn using the *Snn1* genomic sequence. BLAST searches revealed that the two copies were located 123,231 bp and 119,650 bp apart from each other in the Jagger and Norin61 genomes, respectively. Both Jagger and Norin61 were insensitive to SnTox1 despite having two copies of *Snn1*. However, sequence analyses showed that they both had the mutation A2187C in *Snn1-B2* and a deletion at the nucleotide position C2567 in *Snn1-B1*. cDNA sequencing indicated that only *Snn1-B1* was expressed in both lines. BLAST searches of other sequenced wheat lines revealed that ArinaLrFor, Julius, CDC Landmark, Mace, and SY Mattis were insensitive to SnTox1 because they carried neither copy of *Snn1*. Both the hard red spring wheat (HRSW) variety Glenn and the soft white wheat variety Fielder possessed *Snn1-B1* with multiple mutations within the gene and were insensitive to SnTox1 (Supplemental Table S3). The HRSW varieties Lancer and Rollag were also insensitive and possessed the critical mutation G1132C in *Snn1-B2* (Fig. 2).

### Nucleotide diversity and haplotype analysis

Analysis of the coding region of *Snn1-B1* revealed 58 SNPs that corresponded to 37 amino acid changes. These polymorphisms represented 22 coding sequence haplotypes (16 insensitive and 6 sensitive) based on the nucleotide sequences that corresponded to 16 amino acid isoforms based on the predicted amino acid sequences (Fig. 3, Supplemental Fig. S1, Supplemental Table S2). However, only six coding sequence haplotypes (3 insensitive and 3 sensitive) corresponding to five amino acid isoforms were identified for *Snn1-B2* (Fig. 4, Supplemental Fig. S2, Supplemental Table S2). A comparative analysis of the complete genomic region of *Snn1* genes among all cultivated wheat lines revealed high levels of nucleotide diversity in *Snn1-B1* compared to *Snn1-B2* (Table 1). The overall nucleotide diversity (π) of *Snn1-B1* was 0.0015, whereas that of *Snn1-B2* was 0.00066. In addition, the genetic variabilities of *Snn1* genes were evaluated using the Tajima’s *D* (Tajima 1989) and the Fu and Li neutrality tests (Fu and Li 1993) to determine if the target gene sequences fit the neutrality model of evolution. According to the results, Fu and Li’s *D* and *F* values for the full genomic region of *Snn1-B1* among all cultivated lines were significantly negative (Table 1). All three neutrality test statistics including Tajima’s *D* and Fu and Li’s tests were significantly negative in *Snn1-B2* among the tetraploid wheat varieties (Table 2). Although Tajima’s *D* and Fu and Li’s *F* test statistics were also negative for *Snn1-B2* among the hexaploid wheat varieties, those values were not significant (*P* > 0.10). Fu and Li’s *D* test statistic had a positive value (0.14046) for *Snn1-B2* among the hexaploid wheat lines.

**Figure 3.**
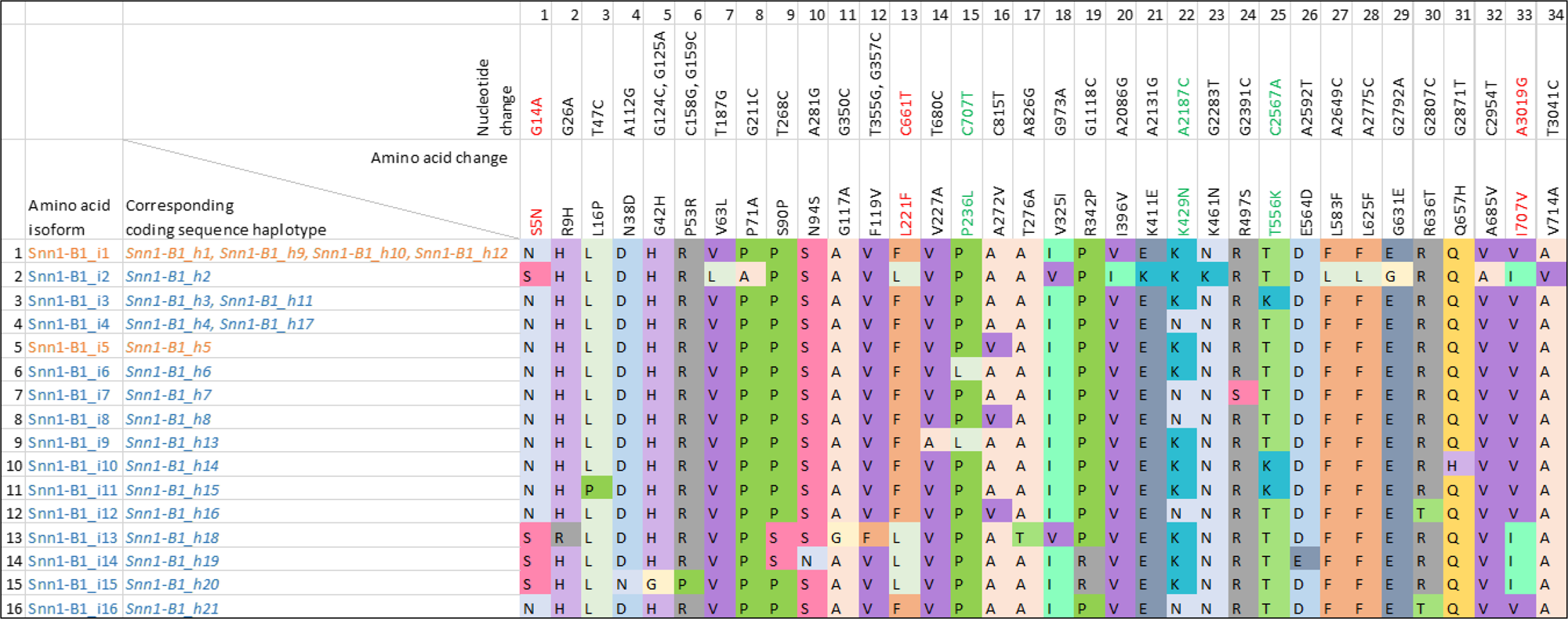
Isoforms identified based on the deduced amino acid sequences of Snn1-B1. Red font: change from insensitive to sensitive; Green font: change from sensitive to insensitive. Orange font: Sensitive haplotypes/isoforms; Blue font: Insensitive haplotypes/isoforms.

**Figure 4.**
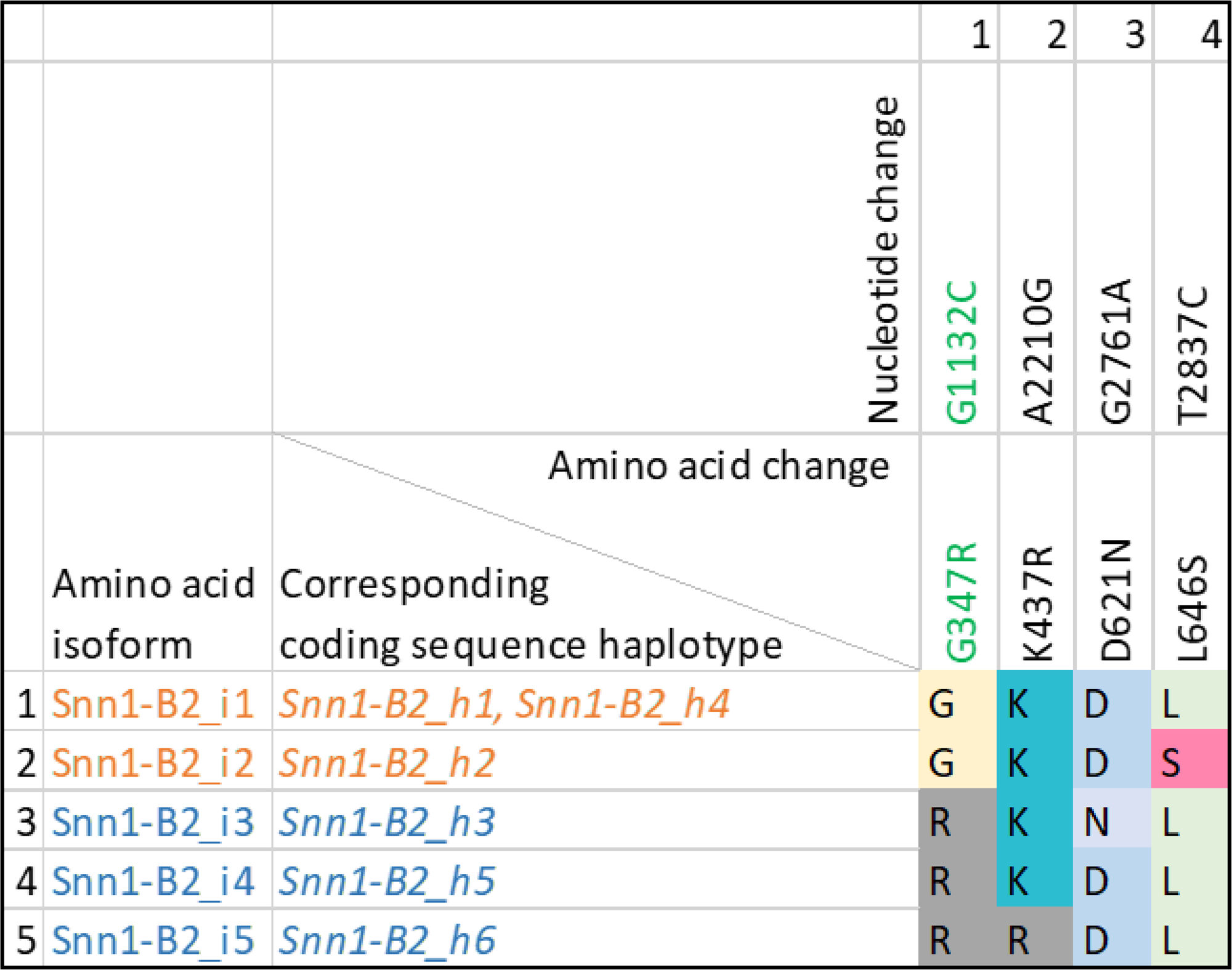
Isoforms identified based on the deduced amino acid sequences of Snn1-B2. Green font: change from sensitive to insensitive. Orange font: Sensitive haplotypes/isoforms; Blue font: Less sensitive haplotypes/isoforms.

**Table 1.** Comparison of nucleotide diversity and neutrality test results of the *Snn1-B1* and *Snn1-B2* genes among all domesticated wheat (tetraploid plus hexaploid) accessions.

| | Number of sites (bp) | $\pi$ | k | $\theta_w$ | Hd | Tajima's *D* | FLD | FLF |
| --- | --- | --- | --- | --- | --- | --- | --- | --- |
| *Snn1-B1* Full gene | 3045 | 0.00150 | 4.57620 | 0.00275 | 0.951 | -1.52547 | -4.14963** | -3.81043** |
| Exon 1 | 991 | 0.00129 | 1.27866 | 0.00156 | 0.734 | -0.44747 | -1.12427 | -1.06559 |
| Intron 1 | 93 | 0.00203 | 0.18868 | 0.01185 | 0.075 | -1.98426* | -3.76872** | -3.75856** |
| Exon 2 | 165 | 0 | N/A | N/A | N/A | N/A | 0 | 0 |
| Intron 2 | 807 | 0.00277 | 2.22932 | 0.00547 | 0.803 | -1.56178 | -4.33536** | -3.99802** |
| Exon 3 | 989 | 0.00089 | 0.87954 | 0.00134 | 0.567 | -0.84241 | -1.43487 | -1.46266 |
| *Snn1-B2* Full gene | 3045 | 0.00066 | 2.00877 | 0.00071 | 0.6 | -0.20579 | -0.50099 | -0.47533 |
| Exon 1 | 991 | 0.00051 | 0.50125 | 0.00022 | 0.501 | 1.70448 | 0.5322 | 1.01076 |
| Intron 1 | 93 | 0 | N/A | N/A | N/A | N/A | 0 | 0 |
| Exon 2 | 165 | 0.00292 | 0.48246 | 0.00131 | 0.482 | 1.59183 | 0.5322 | 0.97286 |
| Intron 2 | 807 | 0.00054 | 0.43734 | 0.00107 | 0.135 | -1.09387 | 0.99173 | 0.40334 |
| Exon 3 | 989 | 0.00059 | 0.58772 | 0.00088 | 0.529 | -0.71138 | -2.3781 | -2.1762 |

$\pi$ Nucleotide diversity per site; k Average number of nucleotide differences; $\theta_w$ Theta per site; Hd Haplotype gene diversity; FLD Fu and Li's *D* test statistic; FLF Fu and Li's *F* test statistic; \* Significant at $P < 0.05$ ; \*\*Significant at $P < 0.02$ ; N/A Not applicable.

**Table 2.** Nucleotide diversity and neutrality test of different regions along the *Snn1-B2* gene within tetraploid and hexaploid domesticated wheat accessions.

| | | Number of sites (bp) | $\pi$ | k | $\theta_w$ | Hd | Tajima's *D* | FLD | FLF |
| --- | --- | --- | --- | --- | --- | --- | --- | --- | --- |
| Hexaploid wheat | Full gene | 3045 | 0.00039 | 1.19203 | 0.0007 | 0.377 | -1.42317 | 0.14046 | -0.37036 |
|  | Exon 1 | 991 | 0 | N/A | N/A | N/A | N/A | 0 | 0 |
|  | Intron 1 | 93 | 0 | N/A | N/A | N/A | N/A | 0 | 0 |
|  | Exon 2 | 165 | 0.00097 | 0.15942 | 0.00162 | 0.159 | -0.68111 | 0.62273 | 0.3105 |
|  | Intron 2 | 807 | 0.00088 | 0.70652 | 0.00133 | 0.236 | -0.93098 | 1.08443 | 0.5923 |
|  | Exon 3 | 989 | 0.00033 | 0.32609 | 0.00081 | 0.308 | -1.49431 | -1.35733 | -1.61079 |
| Tetraploid wheat | Full gene | 3045 | 0.00012 | 0.36364 | 0.00049 | 0.119 | -2.10255* | -3.54262** | -3.62445** |
|  | Exon 1 | 991 | 0.00006 | 0.06061 | 0.00025 | 0.061 | -1.14014 | -1.71333 | -1.78961 |
|  | Intron 1 | 93 | 0 | N/A | N/A | N/A | N/A | 0 | 0 |
|  | Exon 2 | 165 | 0 | N/A | N/A | N/A | N/A | 0 | 0 |
|  | Intron 2 | 807 | 0.0003 | 0.24242 | 0.00122 | 0.061 | -1.88809* | -3.0747* | -3.16649* |
|  | Exon 3 | 989 | 0.00006 | 0.06061 | 0.00025 | 0.061 | -1.14014 | -1.71333 | -1.78961 |

$\pi$ Nucleotide diversity per site; k Average number of nucleotide differences; $\theta_w$ Theta per site; Hd Haplotype gene diversity; FLD Fu and Li's *D* test statistic; FLF Fu and Li's *F* test statistic; \* Significant at $P < 0.05$ ; \*\*Significant at $P < 0.02$ ; N/A Not applicable.

### Protein modeling and structures

According to the predicted protein structures of Snn1-B1 and Snn1-B2, the protein kinase domain consisted of both alpha helices and beta sheets, whereas the GUB_WAK and EGF_CA domains were made with only beta sheets (Fig. 2). The transmembrane domain consisted of one alpha helix. The overall predicted local-distance difference test (pLDDT) scores of Snn1-B1 and Snn1-B2 wild-type protein structures were high at 83 and 85.4, respectively (Supplemental Fig. S3). According to predicted aligned error (PAE) scores of the protein structures, the relative positions of transmembrane domain and S/TPK domain have a high confidence (Supplemental Fig. S4). The relative predicted positions of GUB_WAK and EGF_CA domains also have a high confidence. However, the relative positions of all domains within a predicted structure have a low overall confidence based on PAE scores. The predicted protein structures of Snn1-B1 and Snn1-B2 were almost identical to each other (Fig. 2) with a high TM score of 0.96 (Supplemental Table S4) as expected because of the 99.9% amino acid sequence similarity between the two genes. Analyses using ConSurf (Ashkenazy et al. 2016) revealed that the S/TPK domain, which is located within the cytoplasm, is more conserved than the rest of the protein (Fig. 5). The SnTox1-binding region of the Snn1 protein showed relatively lower evolutionary conservation compared to the other extracellular regions of Snn1.

**Figure 5.**
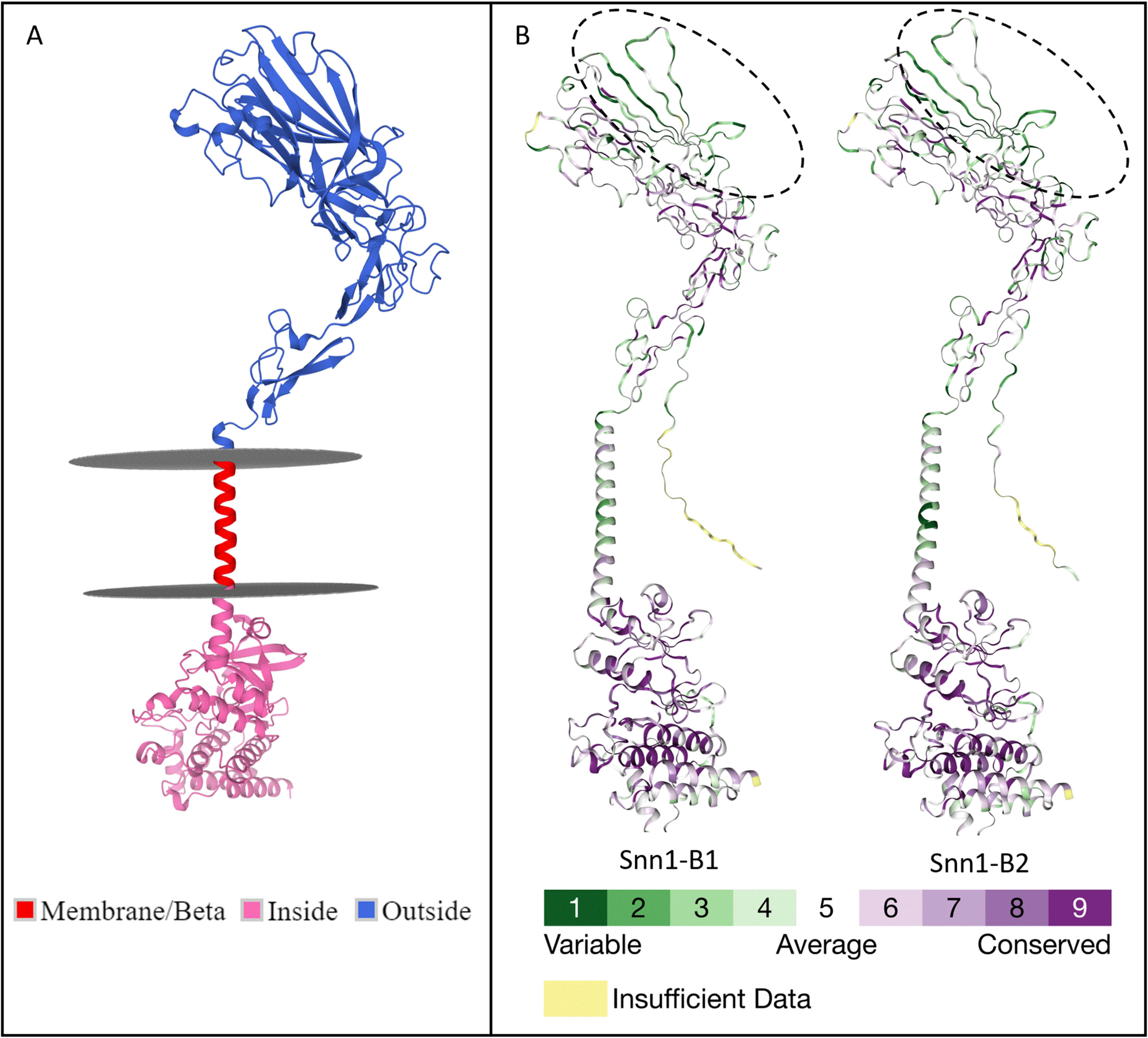
A: Predicted position of the mature Snn1-B1 protein across the cell membrane. B: Evolutionary conservation of Snn1-B1 and Snn1-B2 proteins. Highly conserved residues of the proteins are shown in purple.

### Identification of critical SNPs

According to the phylogenetic analysis, the SnTox1-sensitive lines fell into two major clades with one consisting of *Snn1-B1* and the other consisting of *Snn1-B2* (Fig. 6, Supplemental Fig. S5). These two clades differed by a single missense mutation I43V, which likely occurred upon duplication of *Snn1-B1* to form *Snn1-B2*. In addition, there was another sensitive clade consisting of three lines originated from *Snn1-B1* that was distinct due to the missense mutation A272V. These three clades represented the amino acid isoforms Snn1-B1_i1, Snn1-B2_i1, and Snn1-B1_i5. The durum variety Altar 84 differed from Snn1-B2_i1 by a single missense mutation (L646S) but was still sensitive to SnTox1 (average phenotypic score of 2.60). The insensitive lines formed different clades that were clearly separated from sensitive clades (Fig. 6, Supplemental Fig. S5). The line Botno was considered slightly sensitive to SnTox1 (average phenotypic score of 1.12), but it belonged to a clade with the insensitive landrace Kubanka (average phenotypic score of 0.93). However, the phenotypic scores for these two lines were not significantly different (LSD = 0.5009 at α =0.05).

**Figure 6.**
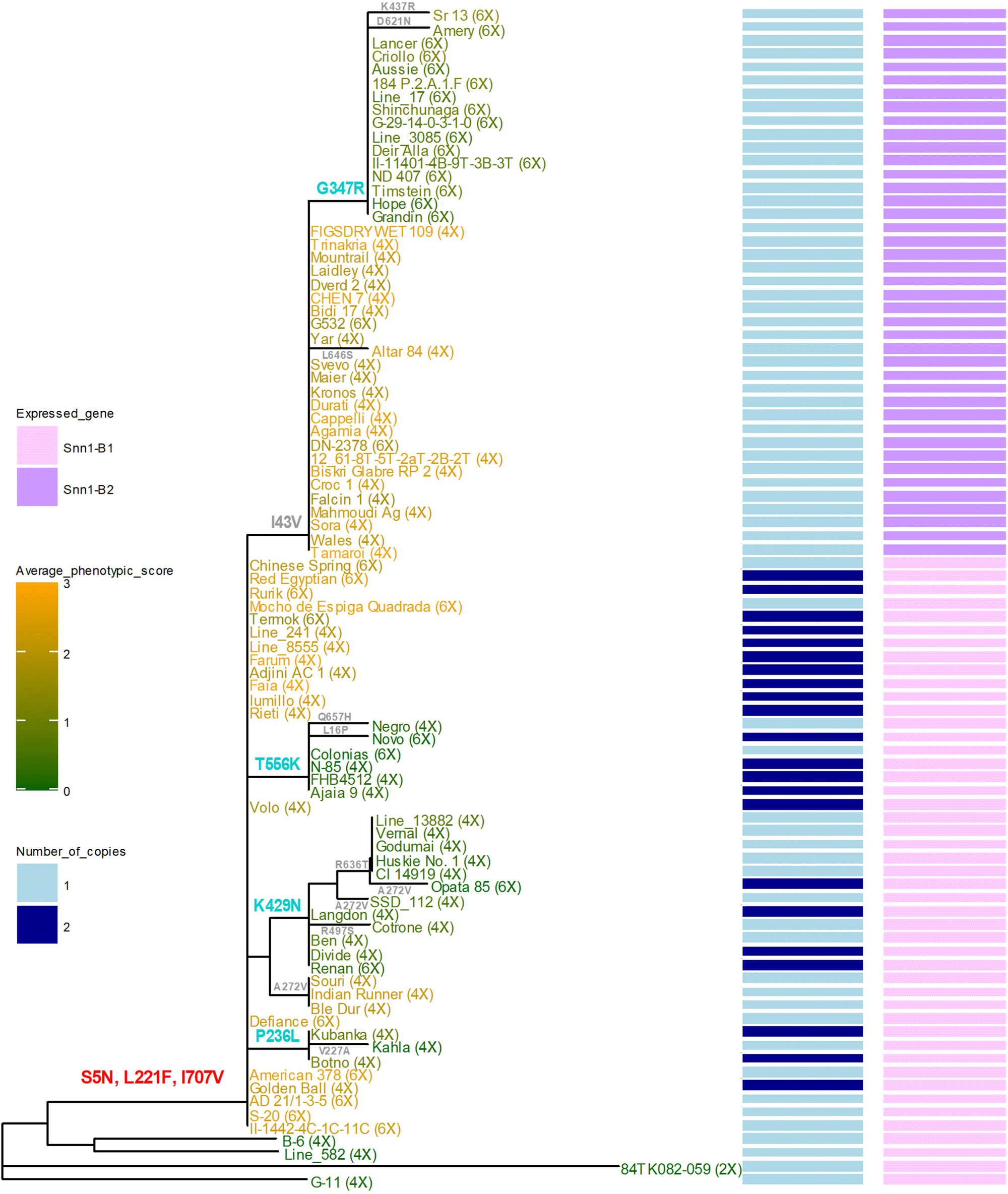
Combined phylogenetic tree of Snn1-B1 and Snn1-B2 based on the deduced amino acid sequences. The color scheme of the heat map is based on the average phenotypic score of each accession. Critical mutations are shown in red (insensitive to sensitive), cyan (sensitive to insensitive) and gray (no change in phenotype) colors.

Phylogenetic analysis of the 114 lines revealed a total of four causal SNPs (C707T, G1132C, A2187C, C2567A) that changed the phenotype from highly sensitive to mostly insensitive (Fig. 6, Supplemental Fig. S5). All four SNPs caused missense mutations that led to amino acid changes P236L, G347R, K429N and T556K, respectively (Figs. 2, 3, 4). Three of these SNPs (C707T, A2187C, C2567A) were found in *Snn1-B1* and caused missense mutations that led to amino acid changes P236L, K429N and T556K, respectively. The fourth SNP (G1132C) occurred in *Snn1-B2* and generated the less sensitive clade G347R in the phylogenetic tree. The SNPs G1132C and A2187C were previously reported by Shi et al. (2016). All four SNPs generate amino acid changes that were expected to be deleterious based on both SIFT (Sorting Intolerant From Tolerant) (Ng and Henikoff 2003) and PROVEAN (Protein Variation Effect Analyzer) (Choi et al. 2012; Choi and Chan 2015) scores (Table 3). The four SNPs occurred in all three exons of the gene showing the importance of all domains for SnTox1 sensitivity.

**Table 3.**
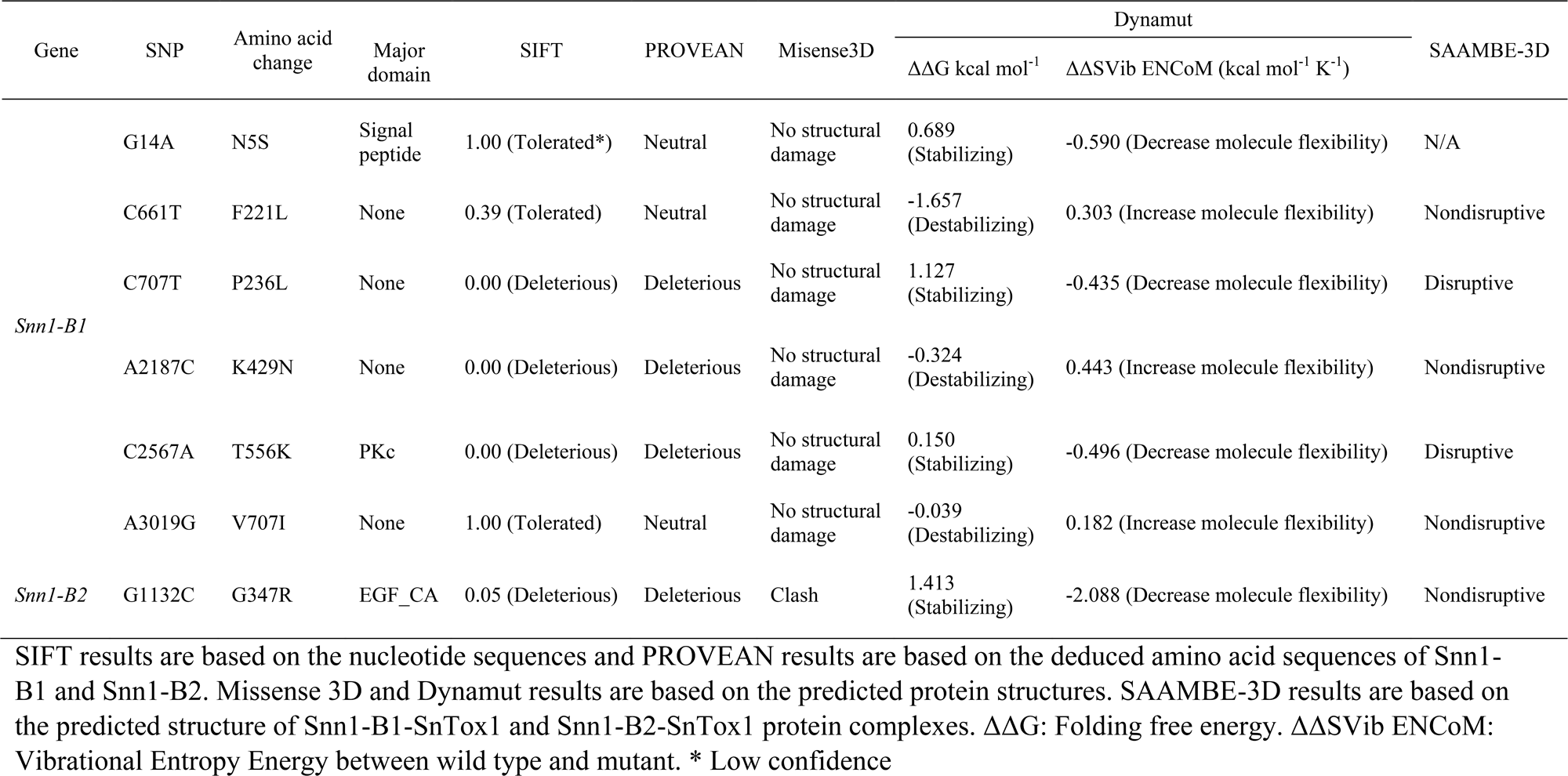
Descriptions and expected impacts of critical mutations identified within *Snn1-B1* and *Snn1-B2* genes.

| Gene | SNP | Amino acid change | Major domain | SIFT | PROVEAN | Misense3D | Dynamut |  | SAAMBE-3D |
| --- | --- | --- | --- | --- | --- | --- | --- | --- | --- |
| | | | | | | | $\Delta\Delta G$ kcal mol<sup>-1</sup> | $\Delta\Delta S_{Vib}$ ENCoM (kcal mol<sup>-1</sup> K<sup>-1</sup>) | |
| *Snn1-B1* | G14A | N5S | Signal peptide | 1.00 (Tolerated*) | Neutral | No structural damage | 0.689 (Stabilizing) | -0.590 (Decrease molecule flexibility) | N/A |
|  | C661T | F221L | None | 0.39 (Tolerated) | Neutral | No structural damage | -1.657 (Destabilizing) | 0.303 (Increase molecule flexibility) | Nondisruptive |
|  | C707T | P236L | None | 0.00 (Deleterious) | Deleterious | No structural damage | 1.127 (Stabilizing) | -0.435 (Decrease molecule flexibility) | Disruptive |
|  | A2187C | K429N | None | 0.00 (Deleterious) | Deleterious | No structural damage | -0.324 (Destabilizing) | 0.443 (Increase molecule flexibility) | Nondisruptive |
|  | C2567A | T556K | PKc | 0.00 (Deleterious) | Deleterious | No structural damage | 0.150 (Stabilizing) | -0.496 (Decrease molecule flexibility) | Disruptive |
|  | A3019G | V707I | None | 1.00 (Tolerated) | Neutral | No structural damage | -0.039 (Destabilizing) | 0.182 (Increase molecule flexibility) | Nondisruptive |
| *Snn1-B2* | G1132C | G347R | EGF_CA | 0.05 (Deleterious) | Deleterious | Clash | 1.413 (Stabilizing) | -2.088 (Decrease molecule flexibility) | Nondisruptive |

SIFT results are based on the nucleotide sequences and PROVEAN results are based on the deduced amino acid sequences of Snn1- B1 and Snn1-B2. Missense 3D and Dynamut results are based on the predicted protein structures. SAAMBE-3D results are based on the predicted structure of Snn1-B1-SnTox1 and Snn1-B2-SnTox1 protein complexes. $\Delta\Delta G$ : Folding free energy. $\Delta\Delta S_{Vib}$ ENCoM: Vibrational Entropy Energy between wild type and mutant. \* Low confidence

In addition, three SNPs were identified as the possible evolutionary causal SNPs that changed the primitive recessive allele of *Snn1-B1* for insensitivity into an allele that caused a sensitive phenotype (Figs. 6, 2, Supplemental Fig. S5). Located at nucleotide positions G14A, C661T and A3019G, all three SNPs caused missense mutations resulting in amino acid changes S5N, L221F and I707V, respectively. It is possible that one or more of these three mutations transformed the insensitive allele in *T. turgidum* ssp*. dicoccoides* to the sensitive *Snn1-B1* allele in durum and common wheat. Locations of the identified critical mutations are shown in Figure 2.

According to DynaMut (Rodrigues et al. 2018), the mutations F221L, K429N and V707I destabilize the Snn1-B1 protein, and the mutations N5S, P236L, T556K, and G347R decrease the flexibility of it (Table 3). Analyses of the impacts of the identified mutations on predicted Snn1-B1-SnTox1 (Fig. 7) and Snn1-B2-SnTox1 protein complexes using SAAMBE-3D (Pahari et al. 2020) revealed that the mutations P236L and T556K can disrupt the complex (Table 3). Analyses of the predicted protein complexes using PDBePISA (Krissinel and Henrick 2007) revealed that F221L is located within the region where Snn1 binds with SnTox1 (Fig. 7). According to Missense3D (Ittisoponpisan et al. 2019), the mutation G347R causes a clash on the Snn1-B2 protein with a local score of 37.41, which is an increase of 22.75 in the clash score compared to the wild-type structure (Fig. 8). No significant change in the secondary structures were observed between wild-type and mutant proteins (Supplemental Fig. S6).

**Figure 7.**
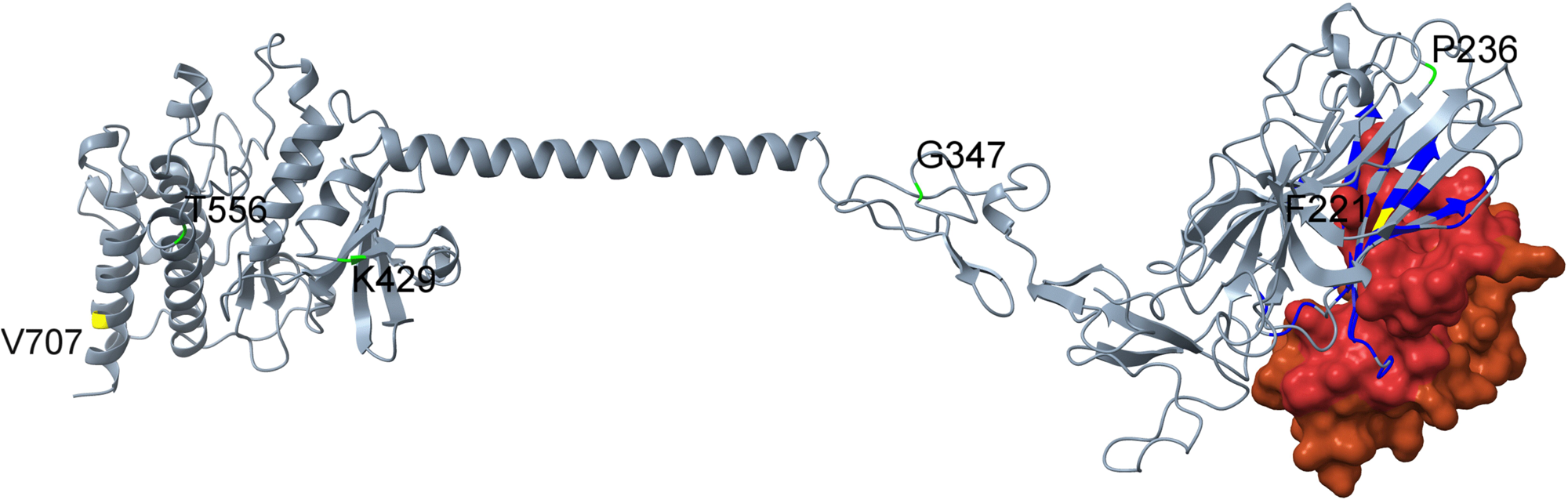
Snn1-SnTox1 protein complex. Gray: Snn1; Brown: SnTox1. Interfacing residues of Snn1 protein are shown in blue and those of SnTox1 protein are shown in red. The amino acid residues associated with the formation of the SnTox1-sensitive Snn1-B1 from a primitive wild wheat are shown in yellow. Amino acids associated with the change in phenotype from sensitive to insensitive are shown in green.

**Figure 8.**
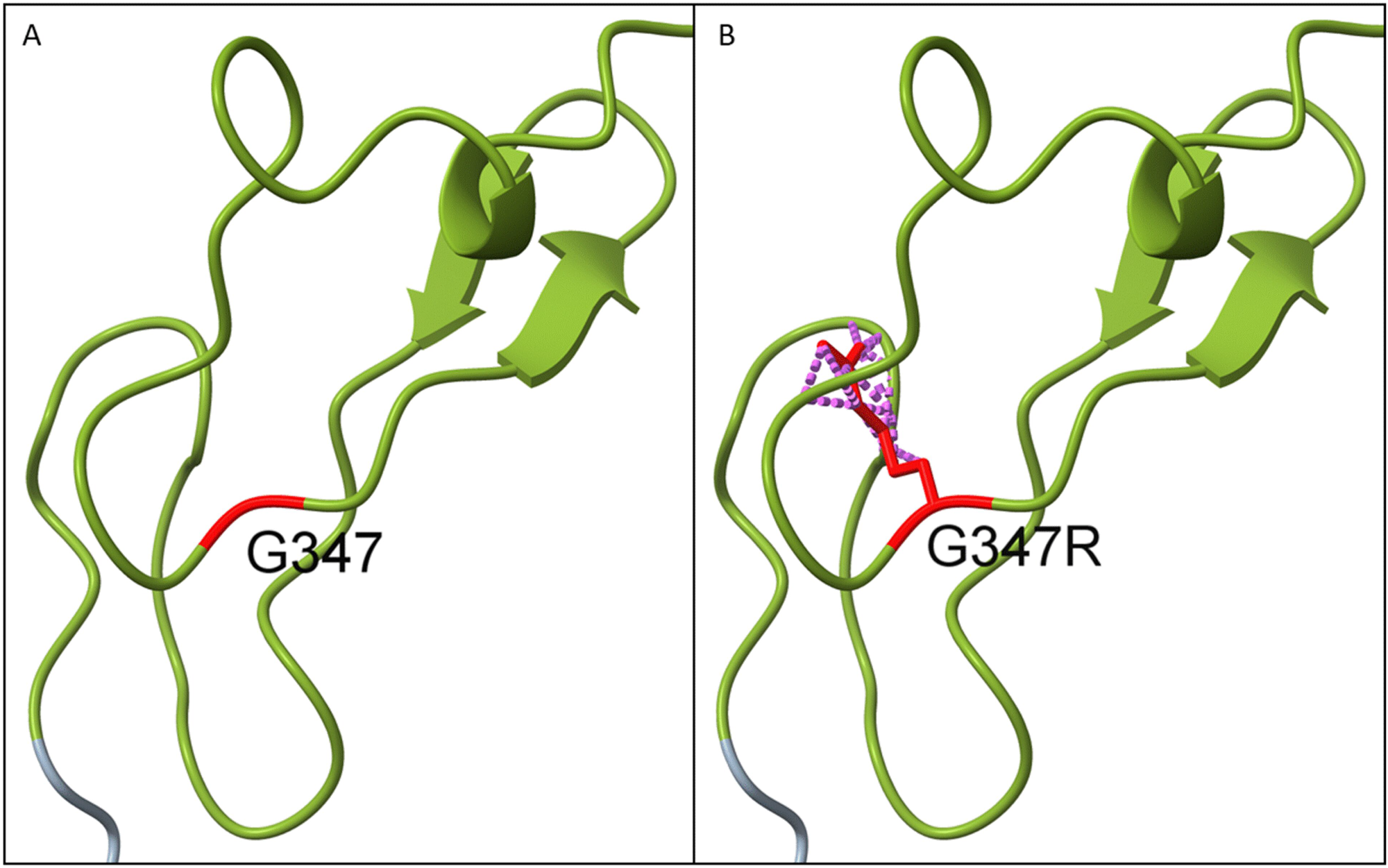
An illustration of the clashes caused by the mutation G347R in Snn1-B2. Residues of interest are highlighted in red and clashing interactions are shown in purple.

The presence of a 4 bp deletion in exon 1 (*Snn1-B1*_584_587del) was observed in five HRSW accessions (PI 94555, PI 180617, PI 168789, CItr 3008, PI 163428) and spelt wheat (PI 190962) carrying *Snn1-B1* (Supplemental Table S3). All these accessions except PI 180617 also possessed a 2 bp deletion in intron 2 (*Snn1-B1*_1679_1680del). None of these accessions were included in the phylogenetic analysis because *Snn1-B1*_584_587del caused a premature stop codon within the gene.

### The less sensitive clade G347R

The clade G347R originated from *Snn1-B2* and consisted of only hexaploid wheat lines (Fig. 6). This clade was unique from the other clades also because it included both insensitive and slightly sensitive accessions. Therefore, an additional phenotyping experiment was conducted to determine if there were significant differences among the phenotypic scores within this clade. The clade included 22 accessions representing three amino acid isoforms (Snn1-B2_i3, Snn1-B2_i4, Snn1-B2_i5). Among them, 20 accessions belonged to the same amino acid isoform (Snn1-B2_i4) and coding sequence haplotype (*Snn1-B2_h5*), but it had lines with average phenotypic scores that ranged from 0.35 to 1.20. The LSD analysis showed that there were significant differences among the phenotypic reactions to SnTox1 among these lines (Supplemental Table S5).

### Kompetitive Allele-Specific PCR markers

Approximately half of the SnTox1-insensitive wheat lines were so because neither *Snn1-B1* nor *Snn1-B2* were present, i.e. they were null for both *Snn1-B1* and *Snn1-B2*. Therefore, we developed a marker (*KASP_Snn1_Null*) to detect the null alleles of *Snn1-B1* and *Snn1-B2* (Table 4). During multiplex PCR, the primers Snn1_Null_KA.2 and Snn1_Null_KR.2 amplified a fragment that was specific to *Snn1-B1* and *Snn1-B2* (amplicon 1) creating a FAM^TM^ fluorescent signal. As a control, we used primers Q_KB.1 and Q_KR.1 to amplify a fragment of the *Q* gene (amplicon 2) with a HEX^TM^ signal. Both amplicons were amplified in accessions that carried either *Snn1-B1* or *Snn1-B2*, which resulted in a ‘heterozygous’ cluster on the scatter plot. However, only amplicon 2 was amplified in accessions that did not carry either of the *Snn1* genes, which created a homozygous HEX^TM^ cluster that allowed the identification of accessions with null alleles (Fig. 9). In addition, KASP markers were developed for three of the four SNPs responsible for generating SnTox1-insensitive alleles. A detailed description of these markers is available in the supplementary material (Supplemental File S1).

**Figure 9.**
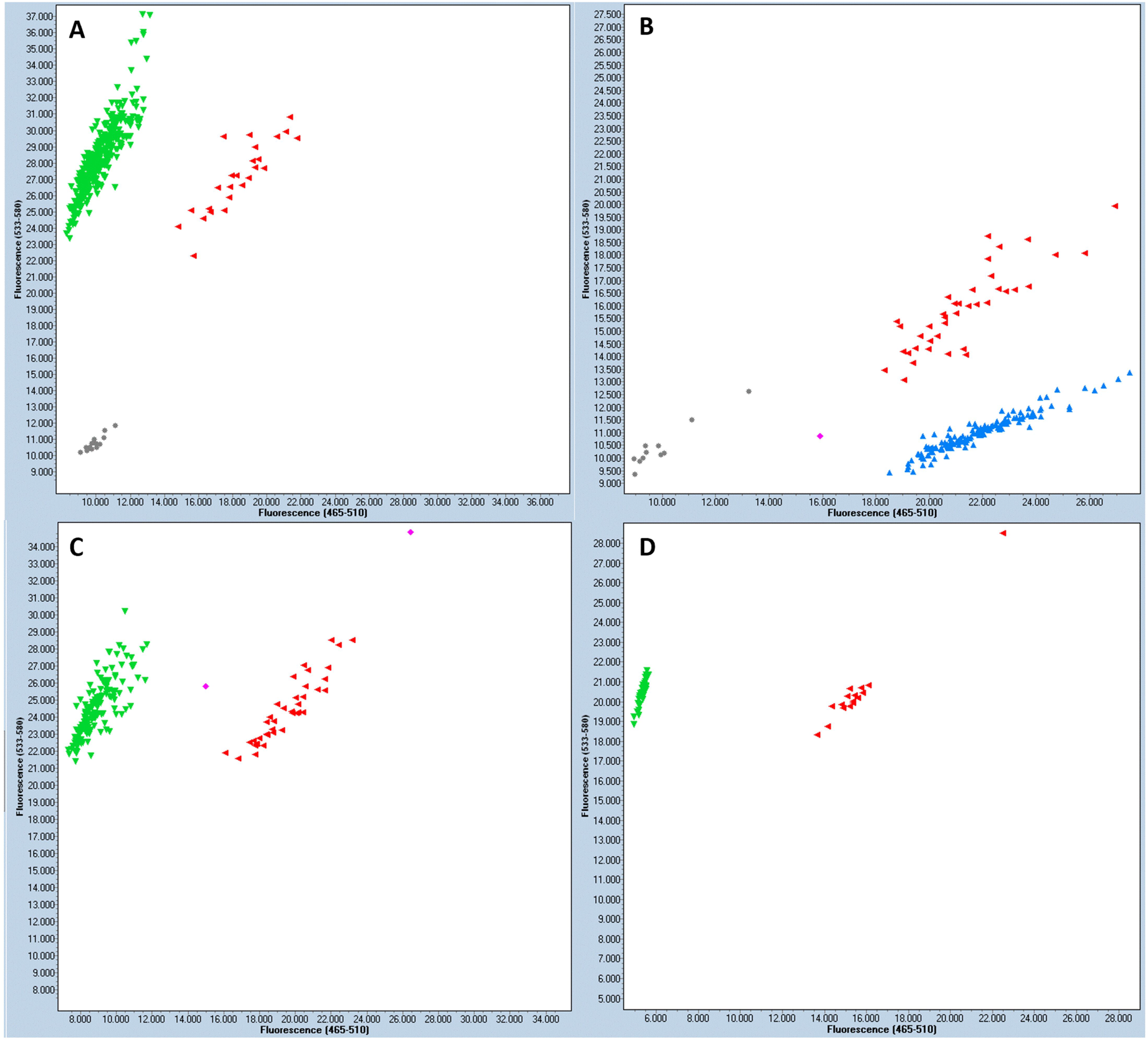
Genotyping scatter plots of the newly developed KASP markers. The X-axes indicate FAM^TM^ fluorescent units (blue) and Y-axes indicate HEX^TM^ fluorescent units (green). Emission of both signals, non-template controls, and unknowns are shown in red, gray, and pink, respectively. A: *KASP_Snn1_C707T*, B: *KASP_Snn1_G1132C*, C: *KASP_Snn1_C2567A*, D: *KASP_Snn1_Null*.

**Table 4.**
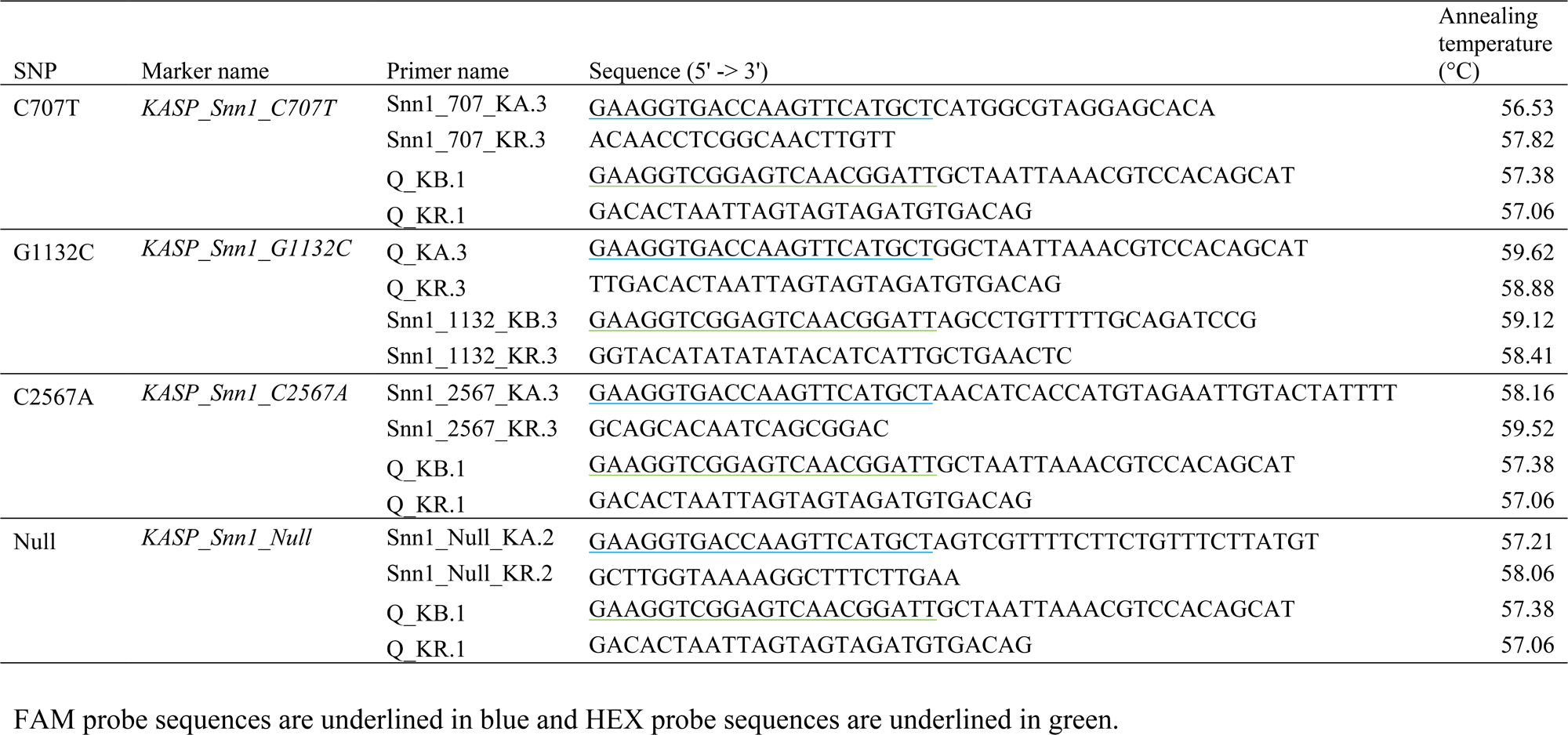
Markers developed to identify lines with critical SNPs.

### Validation and predictability of the markers

The newly developed markers were validated using independent sample sets that included accessions known to carry the corresponding SNPs based on the phylogenetic study as well as negative controls (see Materials and Methods). This validation was done to ensure that each marker consistently predicted the presence or absence of a corresponding SNP in a given accession. According to the results, all four markers accurately predicted the expected genotypes of the accessions used for validation.

Once the markers were validated, their predictabilities of the phenotypes were assessed using 510 accessions of the GDP and 184 HRSW accessions. Here, the association between the genotype of an accession as determined by a marker and its SnTox1 sensitivity score was compared to determine the accuracy of each marker in predicting the phenotype.

According to the KASP assay with the marker *KASP_Snn1_G1132C*, the SNP G1132C was absent in the GDP and is likely specific to hexaploid wheat. The remaining markers *KASP_Snn1_C707T*, *KASP_Snn1_C2567A*, and *KASP_Snn1_Null* together identified 88.3 % of the insensitive lines and 96.9 % of the sensitive lines in the GDP. Collectively, the markers predicted the phenotype of lines in the GDP with an accuracy of 94.7 % (Table 5). Accuracies of the markers *KASP_Snn1_C707T*, *KASP_Snn1_G1132C*, and *KASP_Snn1_C2567A* were lower in the HRSW panel compared to the GDP. The accuracy of *KASP_Snn1_Null* in being able to distinguish null from non-null alleles was 100 % across both panels. However, it was correct in predicting only 54.7 % and 46.9 % of insensitive accessions in the GDP and HRSW panels, respectively, because the remaining percentages of insensitive accessions contained recessive *Snn1* alleles not in the null state. This indicates that null alleles are responsible for roughly half of the insensitive accessions in both panels (Fig. 10).

**Figure 10.**
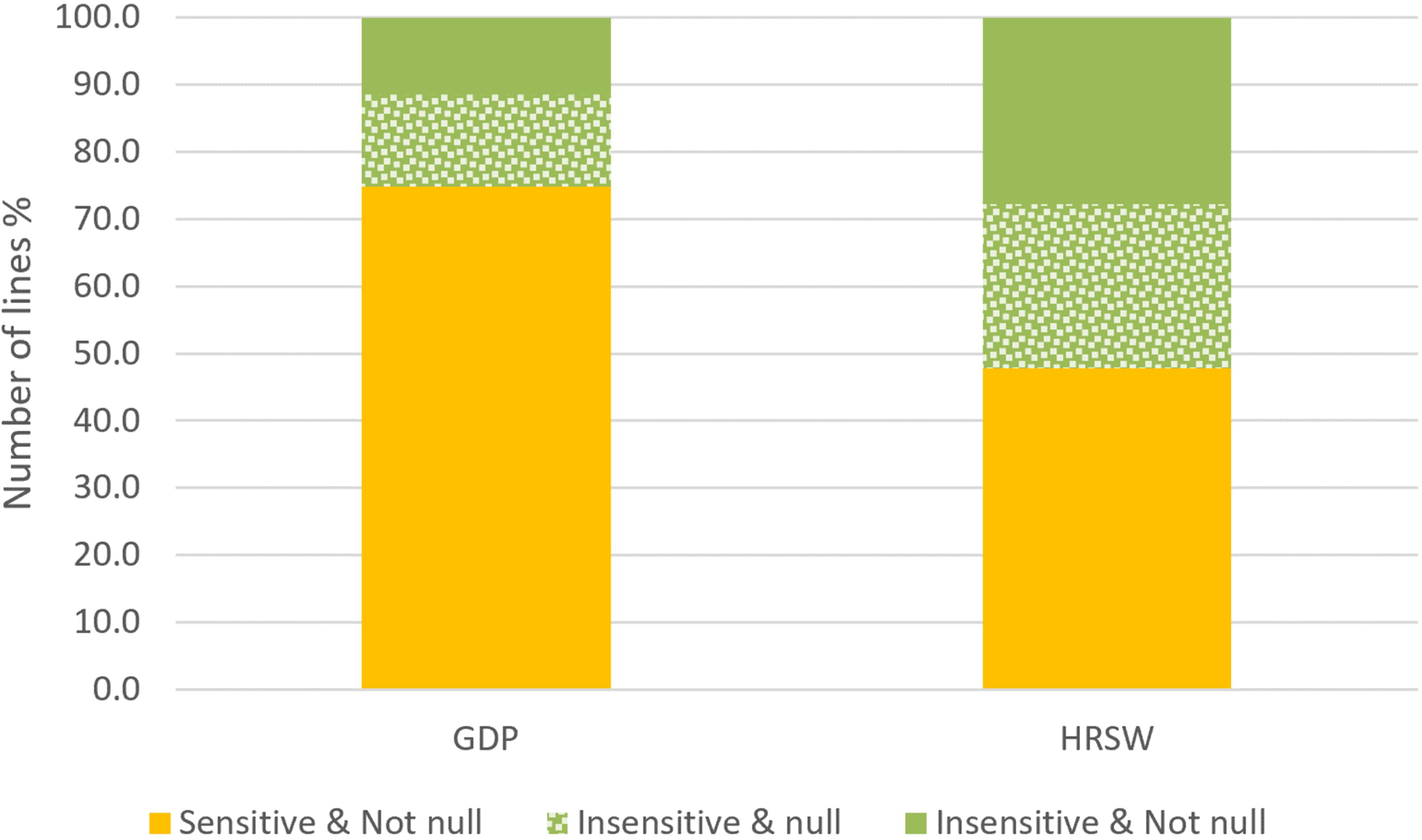
Accuracy of the KASP marker *KASP_Snn1_Null* in the GDP and the HRSW panel. The portion of the panels that were insensitive to SnTox1 is shown in green and the sensitive portion is shown in yellow. The dotted sections represent accessions that were predicted by the marker to contain null alleles. Thus, the marker predicted null genotypes to be insensitive to SnTox1 with 100% accuracy.

**Table 5.** Proportions of the populations explained by newly developed markers and their accuracies on predicting the phenotype.

|  | Accuracy* (%) |  | Proportion explained** (%) |  |
| --- | --- | --- | --- | --- |
|  | GDP | HRSW | GDP | HRSW |
| *KASP_Snn1_C707T* | 82.4 | 5.0 | 21.9 | 1.0 |
| *KASP_Snn1_G1132C* | N/A | 47.6 | 0.0 | 20.8 |
| *KASP_Snn1_C2567A* | 77.4 | 44.7 | 32.0 | 17.7 |
| *KASP_Snn1_Null* | 100.0 | 100.0 | 54.7 | 46.9 |
| All markers collectively predict insensitivity | 90.4 | 65.6 | 88.3 | 85.4 |
| All markers collectively predict sensitivity | 96.1 | 76.3 | 96.9 | 51.1 |
| All markers collectively predict phenotype | 94.7 | 69.0 | 94.7 | 69.0 |
| *KASP_Snn1_Null* predict insensitivity | 100.0 | 100.0 | 54.7 | 46.9 |
| *KASP_Snn1_Null* predict sensitivity | 86.8 | 63.3 | 100.0 | 100.0 |
| *KASP_Snn1_Null* predict phenotype | 88.6 | 72.3 | 88.6 | 71.5 |

\*Percentage of correct predictions made by the marker.
\*\* Percentage of accessions in the panel whose phenotype was correctly predicted by the marker.

## Discussion

### Evolution of *Snn1*

The *Snn1* gene was originally cloned from the common wheat landrace CS, which possessed only one copy, *Snn1-B1* (Shi et al. 2016). At the time, we had hoped that by cloning the gene, the development of functional or diagnostic markers for use in MAE of *Snn1* would be quite straight forward. However, the sequencing and analysis of the diverse set of tetraploid and hexaploid lines described in this research revealed a more complex evolutionary history of *Snn1* including gene duplication, deletion, and multiple causal polymorphisms giving rise to multiple SnTox1-sensitive and -insensitive haplotypes.

Our analysis revealed three SNPs (G14A, C661T and A3019G) that occurred in a primitive insensitive allele of *Snn1-B1* to give rise to the first SnTox1-sensitive allele. The mutations C661T and A3019G are predicted to destabilize the Snn1 protein. A thermodynamically stable, accurately folded, native structure of a protein is important for its functionality (Karplus and Kuriyan 2005, Kumar and Biswas 2019). Among the mutations identified, the SNP C661T caused the highest Gibbs free energy change (ΔΔG) between the wild-type and mutant structures, indicating that it has the highest impact on the stability of the protein. In addition, C661T was predicted to increase the molecular flexibility of the Snn1-B1 protein with the highest vibrational entropy change among the mutations identified. Although both C661T and A3019G do not encode amino acids within any of the conserved functional domains of Snn1-B1, the amino acid encoded by C661T resides in the region where Snn1-B1 binds with SnTox1 as determined by Shi et al. (2016). The mutation G14A is located at the beginning of the gene encoding an amino acid in the signal peptide domain, which is important for translocation of the protein to the cellular membrane. Therefore, it seems that either G14A or C661T is likely the causal mutation that gave rise to the sensitive allele during the domestication of wheat, but further investigation is needed.

Subsequent to the evolutionary formation of the first SnTox1-sensitive allele, three causal mutations (C707T, A2187C and C2567A) occurred within the *Snn1-B1* sensitive allele to give rise to insensitive haplotypes. Also, *Snn1-B1* underwent a gene duplication to give rise to *Snn1-B2*, and it is likely to have happened relatively recently given that the two genes share about 99.9% similarity. After duplication, it appears that lines where *Snn1-B2* retained function underwent loss of *Snn1-B1*. One insensitive haplotype arose through mutation of *Snn1-B2* (G1132C), and it appears this occurred recently and is specific to common (hexaploid) wheat and not present in durum (tetraploid) wheat.

### *Snn1* prevalence and diversity

Of the 510 GDP accessions analyzed in this study, 382 (75 %) were sensitive to SnTox1, whereas only 89 (48 %) of accessions among the 184 HRSW lines analyzed were sensitive. This agrees with Shi et al. (2016) that *Snn1* alleles that confer sensitivity to SnTox1 are more prevalent among durum varieties. A relatively high frequency of sensitive alleles in durum wheat varieties compared to a much lower frequency among hexaploid common wheat varieties suggests that *Snn1* might provide a secondary function that may be more important in durum than common wheat and perhaps compensated for by factor(s) on the D genome in the latter.

Common wheat, durum wheat and cultivated emmer have lower overall nucleotide diversity than wild emmer (Haudry et al. 2007). Initial diversity of wild emmer was reduced by 69% in *T. aestivum* and 84% in *T. durum*. This reduction of nucleotide diversity in wheat is lower than that of most other crop species. The average overall nucleotide diversity of *T. aestivum* A and B genomes was reported to be 0.00059 and 0.0008 in two separate studies (Akhunov et al. 2010, Haudry et al. 2007). The nucleotide diversity of *Snn1-B1* among the cultivated wheat accessions used in this study was higher than the overall diversity at 0.0015. High polymorphism levels in R genes facilitate rapid evolution in response to pathogen evolution (Kuang et al. 2004; Allen et al. 2004; Yang et al. 2008). *Snn1-B2* had lower overall nucleotide diversity compared to *Snn1-B1* indicating that it originated later in the evolutionary history of wheat as suggested earlier.

The theory of neutral molecular evolution (Kimura 1983) states that the majority of DNA polymorphisms are selectively neutral in a population. Therefore, the diversity in a population is due to the introduction or loss of polymorphism by mutations and genetic drift. The finding that the values of two of the three neutrality tests for the entire genomic region of *Snn1-B1* were significantly negative (*P* < 0.05) suggests that this gene may have been under diversifying selection. Although the Tajima’s *D* statistics was also negative for *Snn1-B1*, it was not significant (*P* > 0.10) and thus this observation needs further validation. It was also observed that all three neutrality tests were negative (*P* < 0.05) for *Snn1-B2* among tetraploid wheat compared to hexaploids indicating diversifying selection of *Snn1-B2* among tetraploid wheat. If *Snn1* does in fact possess a secondary function as suggested earlier, it might be the reason for the occurrence of diversifying selection.

### *Snn1* functionality and expression

The degree of necrosis caused by a compatible *Snn1*-SnTox1 interaction is somewhat unique compared to the other host sensitivity gene-NE interactions in the wheat-*P. nodorum* system, because it tends to show a range of sensitivity levels. The SNP G1132C explained some of the variation because it was responsible for the G347R clade consisting of lines with less sensitive reactions. However, the same clade included lines showing phenotypic reactions that were significantly different from each other despite carrying identical *Snn1-B2* alleles. Also, both major sensitive clades had accessions with identical coding sequence haplotypes of *Snn1* but significantly different phenotypic scores. It is possible that variation exists within the promoter regions or other cis/trans factors that could affect gene expression and function, or variation in other genetic factors in the backgrounds of different accessions might influence the phenotype such as allelic variation in genes that operate downstream from the initial *Snn1*-SnTox1 recognition.

Although some of the sensitive lines had both *Snn1-B1* and *Snn1-B2*, cDNA sequencing results showed that only *Snn1-B1* was expressed in these lines. Two SNPs that differentiated the two genes (G127A and A264G) were located in the coding region. The SNP A264G was a synonymous mutation and thus unlikely to have any significant impact on expression of the gene. G127A encoded a missense variant (V43I), which was commonly observed in most of the sensitive accessions with expressed *Snn1-B1* or *Snn1-B2*, suggesting that the SNP G127A was also not responsible for the differences in the expression of the two genes. Shi et al. (2016) showed that *Snn1* in the durum variety Lebsock was not expressed even though it had the same sequence as several other SnTox1-sensitive lines. Hence, the differences in expression observed are likely due to differences in the promoter region or other regulatory factors as suggested earlier.

All four SNPs giving rise to insensitive *Snn1* haplotypes in domesticated wheat caused missense mutations and were expected to have deleterious effects based on both SIFT and PROVEAN scores. The hexaploid wheat-specific mutation G1132C that occurred in *Snn1-B2* triggered a clash alert, which means the new amino acid residue cannot fit without changing the conformation of the protein (Čalyševa and Vihinen 2017). Because the accessions that carry this mutation still show low levels of sensitivity to SnTox1, we speculate that it has less impact on the protein structure compared to the other mutations identified. SAAMBE-3D results suggested that the mutation C707T disrupted the Snn1-B1-SnTox1 protein complex. This aligns with the findings of Shi et al. (2016), which indicated that SnTox1 binds directly to a region between GUB_WAK and EGF_CA domains of Snn1, where C707T is located. It was predicted that A2187C destabilizes the protein while increasing its molecular flexibility. Therefore, it is possible that this mutation may cause the protein to adopt non-native or unstable conformations which in turn disrupt its function.

### *Snn1* functional markers

The results of the phylogenetic analysis led to the identification of causal SNPs, which allowed the development of functional markers that could be used to eliminate the sensitive alleles of *Snn1*. The aim was to develop markers that could identify alleles that belong to any of the three sensitive clades (Fig. 6). KASP markers were chosen for design because of their high-throughput and low assay cost (Semagn et al. 2014). It was also important to develop a marker that could detect *Snn1* null alleles because it was observed that null alleles were responsible for a large portion of SnTox1-insensitive lines in both durum and common wheat, and the primary aim of a breeder is to select for SnTox1-insensitive lines. Although a regular KASP marker could have been developed for this purpose, it would not have been possible to differentiate a real null allele and a possible failed PCR with such a marker. Therefore, a multiplex KASP assay consisting of two primer pairs was designed to detect null alleles. The first primer pair was designed to be specific to *Snn1-B1* and *Snn1-B2* and common to all the *Snn1* haplotypes identified in this study.

Theoretically, the second primer pair could have been developed to amplify any part of the wheat genome except *Snn1-B1* and *Snn1-B2*. However, we designed the second pair to be specific to the *Q* gene because it is present in all cultivated wheat lines and progenitors (Faris and Gill 2002). Amplification by this primer pair acts as a control that indicates a successful PCR. The marker *KASP_Snn1_Null* was 100 % accurate in predicting whether a line was null for *Snn1*. However, the caveat here is that the non-null allele class includes both SnTox1-sensitive lines and lines that carry recessive alleles for SnTox1 insensitivity, i.e. *KASP_Snn1_Null* is not able to distinguish between sensitive and insensitive *Snn1* alleles, but only presence or absence of the gene.

Attempts to develop regular KASP markers that would detect the SNPs C707T, G1132C, A2187C, and C2567A, while creating distinct scatter plots were unsuccessful (data not shown), which was the reason for designing four-primer multiplex KASP markers for these SNPs. The newly developed markers (*KASP_Snn1_C707T*, *KASP_Snn1_G1132C*, *KASP_Snn1_C2567A*), can identify an accession as insensitive if it carries the relevant mutation in either *Snn1-B1*, *Snn1-B2,* or both (in both homozygous and heterozygous state). However, it can also mistakenly classify some of the sensitive accessions carrying both genes as insensitive if only one of the genes has the target mutation. This could be the reason for the low accuracy of these markers in the HRSW panel. However, the accuracy of these markers was high in the durum wheat panel, and the usage of them along with *KASP_Snn1_Null* increased the prediction of insensitive lines from 54.7 % to 88.3 % in the GDP.

The ultimate goal of a breeding program is to develop varieties that are high yielding with good quality. Obtaining resistance to diseases is necessary to achieve that goal, and SNB is an economically important disease of durum and common wheat that must be considered. SNB resistance can be at least partially achieved through removal of *Snn1* alleles to gain insensitivity to SnTox1. Although we developed multiple markers in this study, we recommend that breeders use the marker *KASP_Snn1_Null* because it is 100 % accurate in identifying lines lacking *Snn1* in both durum and hexaploid wheat, making it the most effective marker available for MAE of *Snn1* and thus the selection of SnTox1-insensitive lines. However, the caveat to using only *KASP_Snn1_Null* is that approximately half of SnTox1-insensitive lines would be eliminated along with the sensitive lines. If this would be undesirable for a breeder, they could combine the use of *KASP_Snn1_Null* along with the other KASP markers developed here to obtain additional information regarding lines containing sensitive vs insensitive alleles in the non-null state. In addition, the *Q*-gene-specific primers designed in this study can be used in the future to develop a null marker for any gene of interest in wheat in combination with two gene-specific primers.

## Materials and methods

### Plant materials

A total of 114 wheat accessions including 43 accessions of common wheat (*T. aestivum* ssp. *aestivum*), 55 accessions of durum wheat (*T. turgidum* ssp*. durum*), two accessions of makha wheat (*T. aestivum* ssp*. macha* Dekapr. et MenAbde, 2*n* = 6*x* = 42, AABBDD), two accessions of club wheat [*T. aestivum* ssp*. compactum* (Host) MacKey, 2*n* = 6*x* = 42, AABBDD], eight accessions of cultivated emmer wheat [*T. turgidum* ssp*. dicoccum* (Shrank) Shubl, 2*n* = 4*x* = 28, AABB], three accessions of wild emmer wheat [*T. turgidum* ssp*. dicoccoides* (Körn. Ex Asch. & Graebner) Aarons, 2*n* = 4*x* = 28, AABB], and one accession of the diploid goatgrass *Aegilops speltoides* ssp*. ligustica* Tausch (2*n* = 2*x* = 14, SS) were used for the phylogenetic study (Supplemental Table S2). Accessions were selected from different parts of the world to increase the diversity of the population.

Validation of three newly developed markers was performed using three common wheat accessions and three durum wheat accessions (Botno, Kahla, Amery, Grandin, Negro, and Novo) as positive controls, with two accessions carrying a specific SNP (see results) per marker. In addition, the hexaploid landrace Chinese Spring (CS) (carrying *Snn1-B1*), the durum variety Kronos (carrying *Snn1-B2*), and the CS 1BS deletion line CS 1BS-18 (Gill et al.1996) were used as negative controls during marker validation. Validation of the marker *KASP_Snn1_Null* was done using the CS nullisomic 1B-tetrasomic 1A (CS N1B-T1A) line and the CS 1BS deletion lines CS 1BS-9 and CS 1BS-18 as positive controls along with CS and Kronos as negative controls. Eight replicates of each control accession were used for PCR.

Two large and diverse panels of wheat were infiltrated with SnTox1 to determine the frequency of SnTox1 sensitivity and to evaluate the efficacy of markers. The first panel was a subset of the global durum panel (GDP), which consisted of 510 durum lines and landraces that represent the majority of global diversity (Mazzucotelli et al. 2020). The second panel consisted of a set of 184 HRSW lines, which was a subset of the National Small Grains Collection panel described by Maccaferri et al. (2015). All lines that were genotyped but not included in the phylogenetic study are listed in Supplemental Table S3.

### Phenotyping and statistical analysis

Phenotyping in this study was done in five separate experiments. For all experiments, two plants were grown in plastic containers measuring ∼ 3.8 cm in diameter, which were arranged in racks containing 98 containers per rack. Plants were given adequate fertilizer and water, and they were grown in a growth chamber at 21 °C. Each container of two plants was considered an experimental unit. A detailed description of each experiment is available in the supplementary material (Supplemental File S1).

SnTox1 was obtained from a *Pichia pastoris* culture expressing SnTox1 as described in Szabo-Hever et al. (2023). All plants were infiltrated with SnTox1 as described in Liu et al. (2004a). At least two plants were infiltrated per accession in each replicate of all experiments. All plants were kept at 21 °C and leaves were evaluated on the fifth day after infiltration.

An expanded scoring scale, which included seven categories (0, 0.5, 1.0, 1.5, 2.0, 2.5, and 3.0), was developed for rating SnTox1 sensitivity levels based on the one defined by Zhang et al. (2011) (Fig. 1). The scores 0 and 3.0 were same as the ones defined by Zhang et al. (2011) where 0 = no visible necrosis or chlorosis and 3.0 = extensive and severe necrosis throughout the entire infiltrated area with complete tissue collapse and shriveling or narrowing of the leaf within the infiltrated region. A score of 1.0 represented mottled chlorosis extending to the boundaries of the infiltrated area and a score of 2.0 was used if the infiltrated area had highly visible chlorosis with no visible necrosis. Scores of 0.5, 1.5 and 2.5 were added to the scale to represent sensitivity levels that were intermediate between the ranges of 0 – 1.0, 1.0 – 2.0 and 2.0 – 3.0, respectively. Lines with scores ≤ 1.0 were considered as insensitive and lines with scores > 1.0 were considered as sensitive.

Bartlett’s test for homogeneity of variances (Snedecor and Cochran 1989) and analysis of variance was conducted using SAS 9.4 (SAS Institute Inc.) Fisher’s protected least significant difference (LSD) was also calculated at an α level of 0.05 to identify significant differences of the phenotypic scores among lines.

### Identification of two copies and sequencing

A CAPS marker designated as *fcp667* was developed targeting SNP G127A to test all lines to be sequenced to identify lines with two copies of the gene (Supplemental Fig. S7). However, this marker was not completely diagnostic for the presence of two copies of *Snn1*, and some additional sequencing was needed to characterize the gene in two-copy accessions. The full-length genomic sequence of *Snn1* was obtained from all accessions. A detailed description of PCR conditions and sequencing is available in the supplementary material (Supplemental File S1).

### Sequence data analysis

Sequence reads were assembled into contigs for each accession, and the predicted amino acid sequences were generated from the contigs using the software Geneious Prime 2021.2.2 (Biomatters Ltd). Multiple sequence alignments were generated using MUSCLE (Edgar 2004) in MEGA11 (Tamura et al. 2021). The phylogenetic tree was constructed using the Neighbor Joining method (Saitou and Nei 1987) with poisson model and pairwise-deletion option in MEGA11. Confidence values for branch nodes were calculated using 1000 bootstraps (Felsenstein 1985). The phylogenetic tree was color coded and annotated using the R package ggtree (Yu et al. 2018). A complete phylogenetic tree with all 114 accessions (Supplemental Fig. S5) as well as a refined tree containing only 88 accessions (Fig. 6) were generated. The refined tree was generated to enhance the visual clarity of the tree by removing accessions with identical phenotypic scores within the same clade. Haplotype analysis, nucleotide diversity analysis (Nei 1987), Tajima, and Fu and Li neutrality tests were performed using DnaSP v6 (Rozas et al. 2017) at a significance level of *P* < 0.05. Functional characterization of Snn1-B1 and Snn1-B2 amino acid sequences were done using InterProScan (Paysan-Lafosse et al. 2022).

Protein structures were predicted using ColabFold v1.5.2-patch (Mirdita et al. 2022) with default settings which made the predictions with AlphaFold2 (Jumper et al. 2021) using MMseqs2 (Steinegger and Söding 2017) and HHsearch (Steinegger et al. 2019). The structures of Snn1-B1-SnTox1 and Snn1-B2-SnTox1 protein complexes were predicted using Alphafold2-multimer_v1 in ColabFold v1.5.2-patch (Evans et al. 2021). Alphafold2 predicted five models for each protein. The model with the highest pLDDT score was chosen for monomers, and the one with the highest pTM score was selected for complexes in downstream analyses. The residues involved in the interfaces of Snn1-B1-SnTox1 and Snn1-B2-SnTox1 protein complexes were analyzed using PDBePISA (Krissinel and Henrick 2007). Impacts of the identified mutations were assessed using SIFT (Ng and Henikoff 2003), PROVEAN (Choi et al. 2012, Choi and Chan 2015), Missense3D (Ittisoponpisan et al. 2019), DynaMut (Rodrigues et al. 2018), and SAAMBE-3D (Pahari et al. 2020). SIFT scores were calculated using the Ensemble Variant Effect Predictor (McLaren et al. 2016). Protein structure alignment between Snn1-B1 and Snn1-B2 as well as wild type and mutant structures were done using the pairwise structure alignment tool with jFATCAT (flexible) algorithm on RCSB PDB (http://www.rcsb.org/) (Berman et al. 2000). Protein secondary structure prediction was done using Polyview-2D (Porollo et al. 2004). The position of the mature Snn1 protein across the cell membrane was predicted and visualized using MembraneFold v 0.0.71 (Gutierrez et al. 2022), and the evolutionary conserved regions of the predicted proteins were identified using ConSurf web server (Ashkenazy et al. 2016). All the protein figures except those that show position of the cell membrane and the evolutionary conserved regions were prepared using UCSF ChimeraX 1.6.1 (Pettersen et al. 2021).

### Marker development

Each KASP assay was designed to be a multiplex PCR (Liu et al. 2010) consisting of two primer pairs: one pair specific to *Snn1*, and the second pair specific to the wheat *Q* gene, which is a domestication gene harbored by all tetraploid and hexaploid wheats that governs spike morphology (Simons et al. 2006). More details of the markers and the PCR conditions are available in the supplementary material (Supplemental File S1).

## Acknowledgments

We would like to thank Mary Osenga, Danielle Holmes, Stephanie McCoy, Cayley Steen, and Megan Overlander for technical assistance.

## Author contributions

JD Faris and SS designed the experiments; TLF and SSX provided valuable resources to conduct the experiments; SS, G Shi, ASH, ZZ, ARPH, KLDR, G Singh, RSN, JD Fielder, and JD Faris conducted the experiments and collected the data; SS, JD Faris and PEM analyzed and interpreted the data; and SS and JD Faris wrote the paper.

## Supplemental data

**Supplemental Figure S1.** Haplotypes identified based on the coding sequence of *Snn1-B1*. Red font: change from insensitive to sensitive, Green font: change from sensitive to insensitive. Orange font: Sensitive haplotypes/isoforms, Blue font: Insensitive haplotypes/isoforms.

**Supplemental Figure S2.** Haplotypes identified based on the coding sequence of Snn1-B2. Red font: change from insensitive to sensitive, Green font: change from sensitive to insensitive. Orange font: Sensitive haplotypes/isoforms, Blue font: Less sensitive haplotypes/isoforms.

**Supplemental Figure S3.** Predicted structures of wild type and mutant Snn1-B1 and Snn1-B2 proteins. Each structure model is colored according to pLDDT scores. A: Snn1-B1_WT; B: Snn1-B2_WT; C: Snn1-B1_N5S; D: Snn1-B1_F221L; E: Snn1-B1_P236L; F: Snn1-B1_K429N; G: Snn1-B1_T556K; H: Snn1-B1_V707I; I: Snn1-B2_G347R.

**Supplemental Figure S4.** Predicted protein structures color coded based on PAE scores. A: Snn1-B1_WT; B: Snn1-B2_WT; C: Snn1-B1_N5S; D: Snn1-B1_F221L; E: Snn1-B1_P236L; F: Snn1-B1_K429N; G: Snn1-B1_T556K; H: Snn1-B1_V707I; I: Snn1-B2_G347R. Relative positions of residues are not confident between different colors.

**Supplemental Figure S5.** Complete combined phylogenetic tree of Snn1-B1 and Snn1-B2 based on the deduced amino acid sequences. The color scheme of the heat map is based on the average phenotypic score of each accession. Critical mutations are shown in red (insensitive to sensitive), cyan (sensitive to insensitive) and gray (no change in phenotype) colors.

**Supplemental Figure S6.** Comparison of the secondary structures between wild type and mutant proteins. Critical amino acids are highlighted in magenta. A: Snn1-B1_WT (top) vs Snn1-B2_WT (bottom); B: Snn1-B1_WT (top) vs Snn1-B1_N5S (bottom); C: Snn1-B1_WT (top) vs Snn1-B1_F221L (bottom); D: Snn1-B1_WT (top) vs Snn1-B1_P236L (bottom); E: Snn1-B2_WT (top) vs Snn1-B2_G347R (bottom); F: Snn1-B1_WT (top) vs Snn1-B1_K429N; G: Snn1-B1_WT (top) vs Snn1-B1_T556K (bottom); H: Snn1-B1_WT (top) vs Snn1-B1_V707I (bottom).

**Supplemental Figure S7.** Restriction enzyme digestion profiles of PCR products amplified from genomic DNA using the CAPs marker *fcp667*. Fragments were separated on a 1.5% agarose gel. Lanes 1, 2 and 3 contained digested PCR products from an accession with only Snn1-B1, an accession with only Snn1-B2, and an accession with both Snn1-B1 and Snn1-B2, respectively.

**Supplemental Table S1**. Bartlett’s test of homogeneity of score variance for the phenotyping experiments.

**Supplemental Table S2**. Wheat lines used in the phylogenetic analysis.

**Supplemental Table S3.** Wheat accessions used in the study carrying Snn1 sequences that were not included in the phylogenetic analysis.

**Supplemental Table S4**. Comparison of predicted protein structures between wild type and mutants, and Snn1-B1 and Snn1-B2 proteins.

**Supplemental Table S5**. Average phenotypic score of the wheat accessions belong to clade G347R.

**Supplemental Table S6**. Nucleotide diversity and neutrality test of different regions along the Snn1-B1 gene among the domesticated wheat accessions.

**Supplemental File S1**. KASP markers and genotyping conditions, phenotyping experiments, and polymerase chain reaction (PCR) and sequencing.

## Data availability

The authors affirm that all data necessary for confirming the conclusions of the article are present within the article, figures, tables and supplementary materials. Previously undescribed sequences have been deposited in GenBank under accession numbers MN908613 through MN908638, MN954887 through MN954929, and OP823498 through OP823546.

